# Ratiometric assays of autophagic flux in zebrafish for analysis of familial Alzheimer’s disease-like mutations

**DOI:** 10.1101/272351

**Authors:** Haowei Jiang, Morgan Newman, Dhanushika Ratnayake, Michael Lardelli

## Abstract

Protein aggregates such as those formed in neurodegenerative diseases can be degraded via autophagy. To assess changes in autophagic flux in zebrafish models of familial Alzheimer’s disease (fAD) mutations, we first developed a transgene, polyQ80-GFP-v2A-GFP, expressing equimolar amounts of aggregating polyQ80-GFP and a free GFP internal control in zebrafish embryos and larvae. This assay detects changes in autophagic flux by comparing the relative strength of polyQ80-GFP and free GFP moiety signals on western immunoblots probed with an antibody detecting GFP. However, the assay’s application is limited by the toxicity of polyQ80-GFP, and because aggregation of this protein may, itself, induce autophagy. To overcome these issues, we subsequently developed a similar ratiometric assay where expression of a GFP-Lc3a-GFP transgene generates initially equimolar amounts of GFP-Lc3a (directed to autophagic degradation) and a free GFP internal control. The sensitivity of this latter assay is reduced by a cellular protease activity that separates Lc3a from GFP-Lc3a, thus contributing to the apparent free GFP signal and somewhat masking decreases in autophagic flux. Nevertheless, the assay demonstrates significantly decreased autophagic flux in zebrafish lacking *presenilin2* gene activity supporting that the Presenilin2 protein, like human PRESENILIN1, plays a role(s) in autophagy. Zebrafish heterozygous for a typical fAD-like, reading-frame-preserving mutation in *psen1* show decreased autophagic flux consistent with observations in mammalian systems. Unexpectedly, a zebrafish model of the only confirmed reading-frame-truncating fAD mutation in a human *PRESENILIN* gene, the K115Efs mutation of human *PSEN2*, shows possibly increased autophagic flux in young zebrafish (larvae).

## Introduction

Autophagic/lysosomal dysfunction is thought to be involved in the neurodegenerative process of Alzheimer’s disease (AD) (1). The endosomal-lysosomal system is a prominent site for the processing of the AMYLOID BETA A4 PRECURSOR PROTEIN (APP) to form the aggregating peptide amyloidβ (Aβ) (2), which is the major component of the amyloid plaques observed in AD brains. Autophagic vacuoles (AVs) are thought to be the major reservoirs of intracellular Aβ, and accumulation of immature AVs has been detected in AD brains, suggesting that the maturation of AVs to lysosomes may be impaired (3, 4). Both full-length APP and β-secretase-cleaved APP are found in such AVs, which are also highly enriched in PRESENILIN (PSEN) proteins (5) (components of the γ-secretase complexes that cleave APP to form Aβ) suggesting that a link may exist between Aβ production and cell survival pathways through activated autophagy in AD. The Aβ accumulation is thought to be induced by the combination of increased autophagy induction and defective clearance of Aβ-generating AVs (1).

Dominant mutations in the *PRESENILIN* (*PSEN*) genes cause the majority of familial, early onset AD (fAD, (6)). We have previously argued that a body of evidence supports that the AD-relevant effect of these mutations may be to alter the activity of PSEN holoproteins rather than the endoproteolysed forms that are active in the γ-secretase complexes that cleave APP (7). Indeed, in 2010 Lee et al. showed that changes in PSEN1 holoprotein function, rather than γ-secretase activity, appear to affect lysosomal function in *PSEN1* fAD mutant human fibroblasts (8). However, the effects of *PSEN* fAD mutations on autophagy are currently debated (9) and it has not yet been shown that the related protein, PSEN2, plays a role in autophagic flux. Thus, the roles of the multifunctional PSEN proteins in cellular function and AD require further investigation.

Non-mammalian animal models including the zebrafish can facilitate AD research through development of rapid, novel assays for cellular processes such as autophagy. An excellent example is the fluorescent protein-based assay published by Kaizuka et al in 2016 that allows qualitative visualization of differences in autophagic flux within cells (10). However, it is also desirable to make simple, quantitative assessments of relative differences in autophagic flux in whole tissues/animals e.g. when these are subjected to different drug treatments or have different genotypes. In this paper we describe the development of two, internally-controlled assays using zebrafish embryos/larvae that allow rapid quantitative comparison of relative autophagic flux using western immunoblotting. We describe exploitation of the superior of these two assays to demonstrate that activity of the zebrafish *PSEN2*-orthologous gene, *psen2*, is required for efficient autophagy. We also investigate changes in autophagic flux in two novel zebrafish models of fAD-like mutations in the human *PSEN* genes. While a typical, reading frame-preserving fAD-like mutation in zebrafish *psen1* significantly decreases autophagic flux, a model of the only known reading frame-truncating *PSEN* fAD mutation does not and may possibly increase autophagic flux in young fish.

## Methods and Materials

### Zebrafish husbandry and animal ethics

All the wild-type and mutant zebrafish were maintained in a recirculated water system. All work with zebrafish was conducted under the auspices of the Animal Ethics Committee of the University of Adelaide.

### Construction of the polyQ80-GFP-v2A-GFP transgene

A DNA sequence coding for polyQ80-GFP-v2A-GFP (Fig 2A and S1 File A) was synthesized by Biomatik Corp. and subsequently ligated into the pT2AL200R150G (Tol2 transposon-based) (S1 File D) gene transfer vector for expression from the ubiquitously-transcribed elongation factor 1 alpha promoter (EF1α-p) (11). The polyQ80-GFP-v2A-GFP transgene codes for two proteins, an 80-residue polyglutamine repeat sequence (polyQ80) fused to the N-terminal of GFP protein as well as a free GFP protein. A viral 2A peptide (v2A) sequence between the two GFP sequences allows their synthesis as separate entities by a “ribosomal-skip” mechanism (12). Thus, translation of polyQ80-GFP-v2A-GFP mRNA gives 1:1 stoichiometric synthesis of polyQ80-GFP and free GFP. PolyQ80-GFP subsequently aggregates and should be degraded by autophagy, while the free GFP remains soluble in the cytosol to act as an internal control.

**Fig 1.**
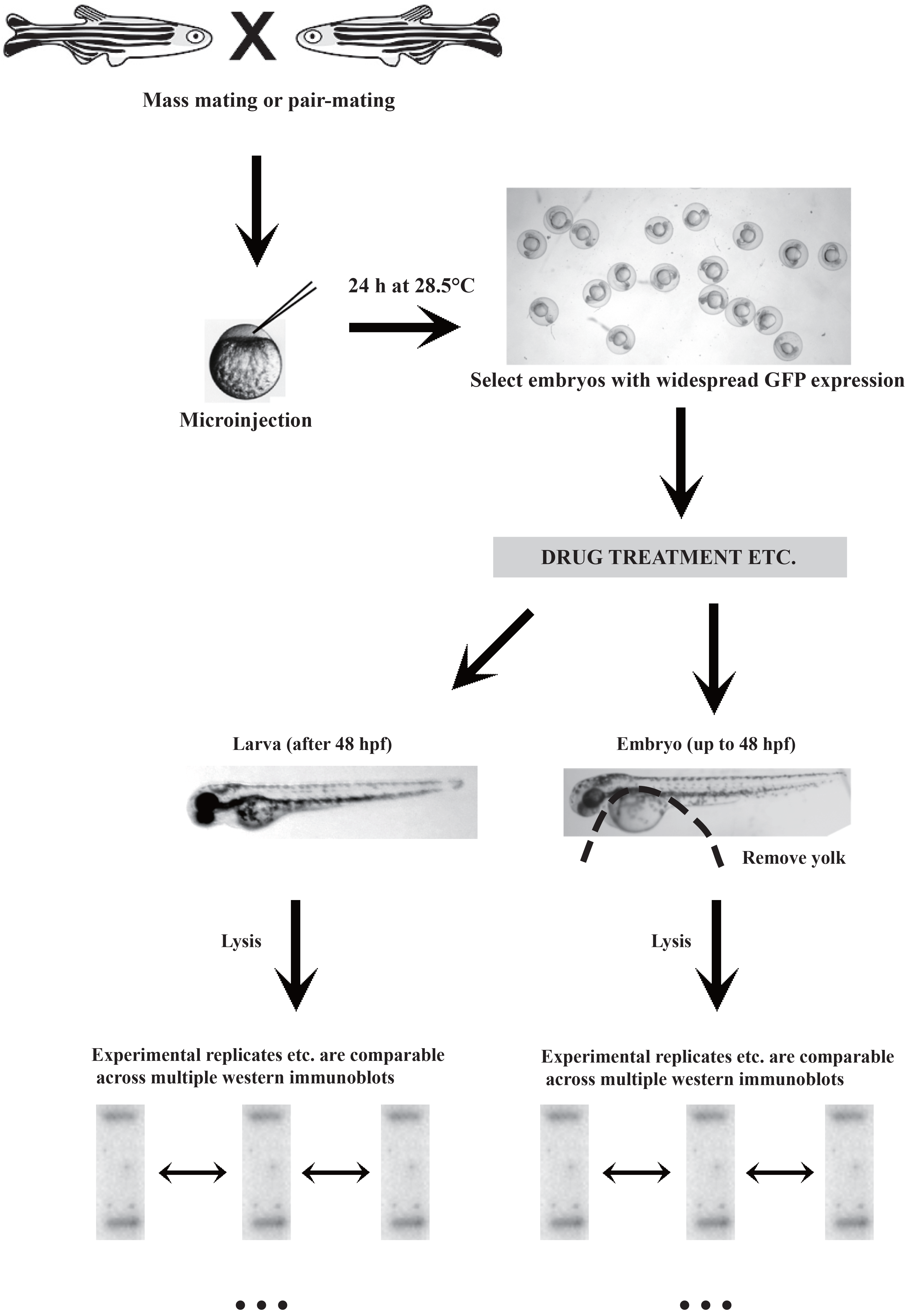
Experimental flow chart.

**Fig 2.**
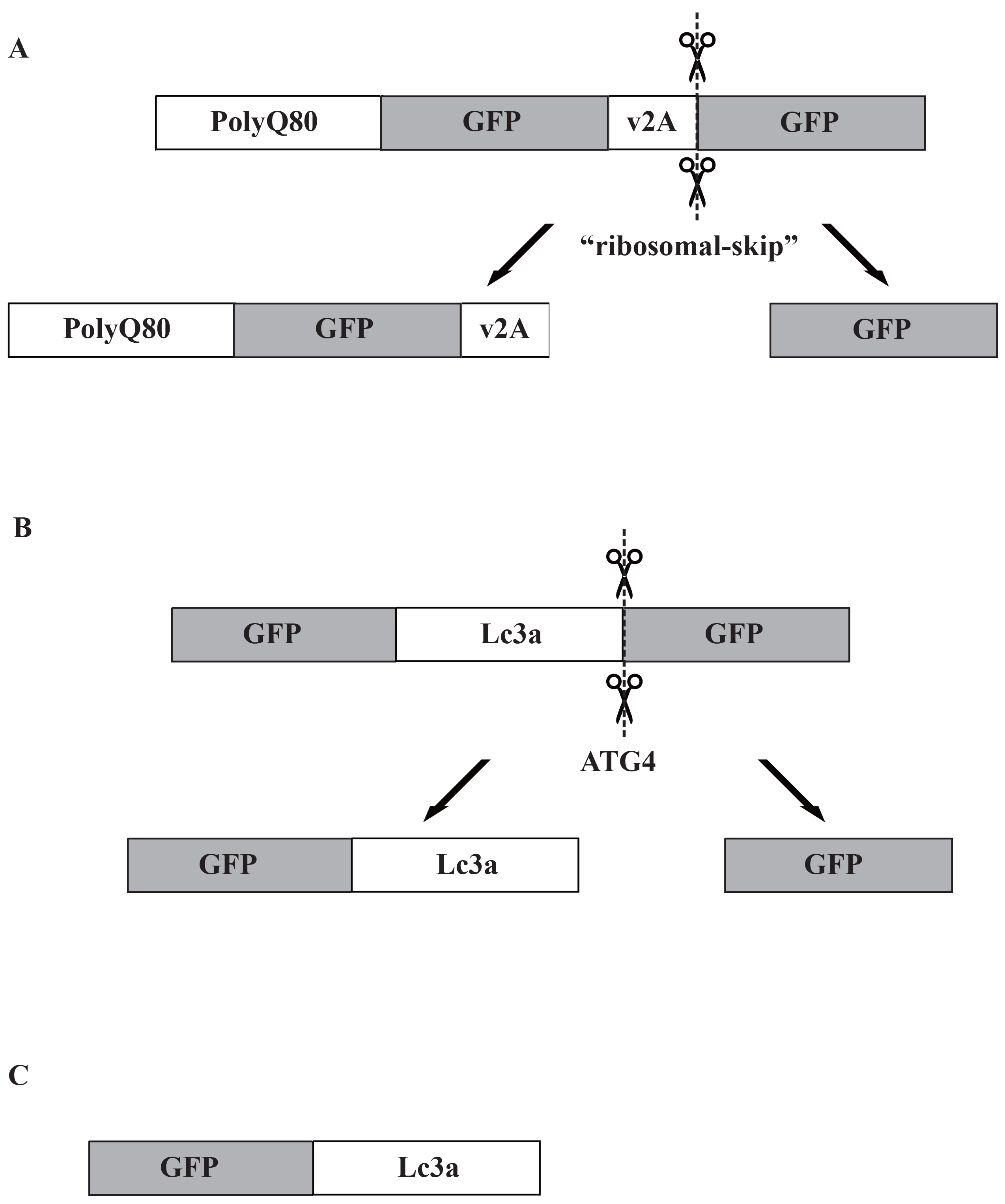
Design of three GFP-based constructs for assay of autophagic flux. (A) Transgene PolyQ80-GFP-v2A-GFP. (B) Transgene GFP-Lc3a-GFP. (C) Transgene GFP-Lc3a.

### Construction of the GPF-Lc3a-GFP transgene

A sequence coding for GFP-Lc3a-GFP (Fig 2B and S1 File B) was synthesized by Biomatik and ligated into the pT2AL200R150G (13) gene transfer vector (S1 File D). This construct encodes a fusion protein where GFP is linked to the N-terminal of zebrafish Lc3a which, at its C-terminal, is linked to an additional GFP. When this transgene is expressed in cells, the most C-terminal glycine residue of Lc3a is cleaved by an endogenous ATG4 family protease, producing equimolar amounts of GFP-Lc3a and free GFP. GFP-Lc3a is conjugated to PE and localizes to autophagosomes (10). The GFP-Lc3a molecules attached to the autophagosomal inner membrane are subsequently degraded after fusion with lysosomes, while those on the outer membrane are deconjugated by Atg4 proteins and recycled back to the cytosol. The free GFP exists in the cytoplasm and functions as an internal control. Relative autophagic activity is measured as changes in the ratio of GFP-Lc3a / free GFP via western immunoblotting.

### Construction of the GFP-Lc3a transgene

The GFP-Lc3a transgene (Fig 2C and S1 File C) in the pT2AL200R150G vector was derived from GFP-Lc3a-GFP by PCR amplification using 5’-phosphorylated primers to exclude the downstream GFP coding sequence followed by ligation to recircularise the plasmid. The sequences of the PCR primers used were 5’-TAGATCGATGATGATCCAGACATGA-3’ and 5’-GCAGCCGAAGGTCTCCT-3’.

### Zebrafish embryos

Three mutations were involved in our research, a putatively null mutation in zebrafish *psen2*, *psen2^S4Ter^*, a model of a typical reading-frame-preserving fAD-type mutation in *psen1*, *psen1^Q96K97del^*, and a model of human *PSEN2^K115Efs^* (14), *psen1^K97Gfs^*. Descriptions of the generation of these mutations were listed in S3 File.

Five different genotypes of zebrafish embryos, Tübingen (TU) wildtype (+/+), putatively *psen2* null heterozygous (*psen2^S4Ter^*/+), putatively *psen2* null homozygous (*psen2^S4Ter^*/*psen2^S4Ter^*), *psen1^Q96K97del^* heterozygous (*psen1^Q96K97del^*/+), and *psen1^K97Gfs^* heterozygous (*psen1^K97Gfs^*/+), were spawned either by mass mating or pair-mating of individuals.

### Microinjection of zebrafish embryos

For the polyQ80-GFP-v2A-GFP assay, zebrafish zygotes were injected with ~5-10 nL of a solution containing 25ng/μL of the polyQ80-GFP-v2A-GFP transgene with 25ng/μL of Tol2 transposase mRNA (11). For the GFP-Lc3a-GFP and GFP-Lc3a assays, zebrafish zygotes were injected with ~5-10 nL of a solution of 50ng/μL of the GFP-Lc3a-GFP or GFP-Lc3a transgenes respectively with 25ng/μL of the Tol2 transposase mRNA. The injected embryos were incubated at 28.5°C in E3 medium (15). At ~24 hours post fertilization (hpf), embryos showing widely distributed GFP expression (as visualized by fluorescence microscopy) were selected for subsequent analysis (Fig 1).

### Rapamycin and Chloroquine treatments

For the PolyQ80-GFP-v2A-GFP assay, GFP-expressing zebrafish embryos from microinjection of polyQ80-GFP-v2A-GFP were separated randomly into three groups at ~30 hpf. Two groups were treated with either 1μM rapamycin (Rapa) (SIGMA, R8781) or 50mM chloroquine (CQ) (SIGMA, C6628) from 30 hpf until 48 hpf, with one group remaining untreated as a control. The embryos were chilled and then lysed for western immunoblotting at ~48 hpf.

For the GPF-Lc3a-GFP assay (or GFP-Lc3a expression analysis) at 48 hpf in +/+ embryos, GFP-expressing zebrafish embryos from microinjection were separated randomly into four groups at ~24 hpf. Three of these groups were treated with rapamycin (1μM) or chloroquine (50μM or 50mM) from 30 hpf until 48 hpf, and the remaining group remained untreated as a control. The embryos were chilled and then lysed immediately for western immunoblotting at ~48 hpf.

For the GPF-Lc3a-GFP assay (or GFP-Lc3a expression analysis) at 96 hpf in +/+ larvae, GFP-expressing zebrafish embryos from microinjection were selected at ~24 hpf and randomly separated into several groups for treatment with rapamycin (1μM) or chloroquine (various concentrations) from 78 hpf until 96 hpf. Larvae were then chilled and lysed immediately for western immunoblotting at ~96 hpf.

### Western immunoblot analyses

48 and 52 hpf-old embryos were firstly dechorionated and deyolked and then placed in sample buffer (2% sodium dodecyl sulphate (SDS), 5% β-mercaptoethanol, 25% v/v glycerol, 0.0625 M Tris–HCl [pH 6.8], and bromophenol blue) (16), heated immediately to 95°C for 10 min and then sonicated (Diagenode Bioruptor UCD-200) in an ice-water bath at high power mode for 10 min before loading onto polyacrylamide gels for electrophoresis (see below). The 96 hpf-old larvae were not deyolked before lysis in sample buffer.

Samples were loaded onto NuPAGE™ 4-12% Bis-Tris Protein Gels (Invitrogen, NP0323BOX), and the separated proteins were subsequently transferred to nitrocellulose membrane (BIO-RAD, 1620115) using the Mini Gel Tank and Blot Module Set (Life technologies, NW2000). The nitrocellulose membranes were subsequently blocked with blocking reagent (Roche, 11921681001) and then probed with the primary antibody, polyclonal anti-GFP goat (ROCKLAND^TM^, 600-101-215), followed by secondary antibody, horseradish peroxidase (HRP) conjugated anti-goat antibody (ROCKLAND^TM^, 605-703-125). Finally, bound antibody was detected by chemiluminescence using SuperSignal™ West Pico PLUS Chemiluminescent Substrate (ThermoFisher, 34580). The ChemiDoc™ MP Imaging System (Bio-Rad) was used to image all the western immunoblots. The intensity of each band from western immunoblots was measured by Image Lab™ Software (Bio-Rad). All these intensity data are presented in S2 File.

### Statistical tests

F-tests were first applied between different groups of data. If the p value of the F-test was >0.05, reflecting no significant difference between the variances of two groups, then a two-tailed t-test assuming equal variances was applied to the two groups. If the p value of the F-test was <0.05, reflecting a significant difference between the variances of two groups, then a two-tailed t-test assuming unequal variances was applied to the two groups.

## Results

### PolyQ80-based autophagy assay in zebrafish

The viral 2A (v2A) system for simultaneous, stoichiometric expression of two peptides from single mRNA transcripts was first applied in zebrafish by Provost et al. (12). The presence of the v2A linker in a coding sequence causes ribosomes to “skip” production of a peptide bond without terminating translation. We sought to exploit this to adapt Ju et al.’s polyQ80-based assay of autophagic flux (described in 2009) to zebrafish. Ju et al expressed fusions to luciferase of aggregating polyQ80 and non-aggregating polyQ21 in separate thigh muscles of mice (17). As a more direct measure of protein concentration we sought to measure by western immunoblotting the relative amounts of a polyQ80-GFP fusion and free GFP when these are translated in a 1:1 ratio from a single mRNA (Figures 1 and 2A). The polyQ80-GFP-v2A-GFP transgene expressing these proteins is carried in a Tol2 transposon vector and introduced into fertilized zebrafish eggs by microinjection together with transposase mRNA (see Materials and Methods, Fig 2A) (Note: We initially developed a polyQ80-GFP-v2A-mCherry-based system but this required immunoblotting against GFP followed by stripping and reprobing the blot to detect mCherry, and the expression ratios thus produced were then only comparable between the samples on individual blots – data not shown.) To allow stable propagation of the polyQ80-GFP-v2A-GFP transgene in *Escherichia coli* (i.e. to avoid recombination between the directly repeated GFP coding sequences) we introduced numerous silent mutations into the degenerate codon positions in the downstream, free GFP coding sequence (see Supplementary Data File 1).

Injection of the polyQ80-GFP-v2A-GFP transgene into fertilized zebrafish eggs results in its widespread insertion into the chromosomes of embryonic cells. Expression of GFP is detectable for weeks afterwards although the transgene does not appear to transmit through the germline (data not shown). Western immunoblotting of embryos (up to 48 hours post fertilization, hpf) or larvae (after 48 hpf) from these injected eggs identified three expected bands of protein: a small fraction of full-length polyQ80-GFP-v2A-GFP, and greater amounts of separated polyQ80-GFP and free GFP proteins (Fig 3A).

**Fig 3.**
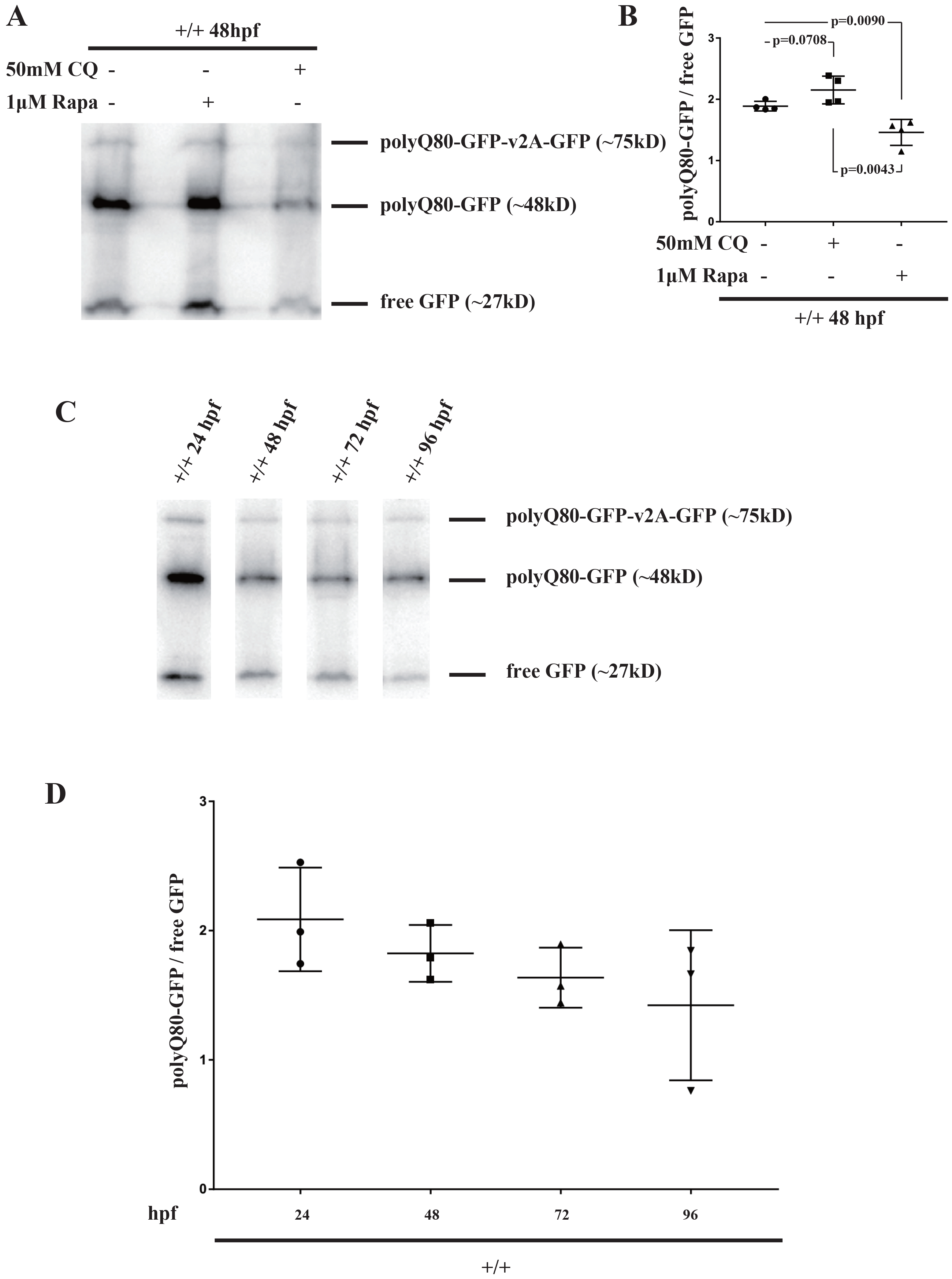
polyQ80-GFP-v2A-GFP assay of autophagy. (A) Western immunoblots from polyQ80-GFP-v2A-GFP-injected +/+ embryos at 48 hpf after treatment with rapamycin (Rapa) or chloroquine (CQ). (B) Ratios (polyQ80-GFP / free GFP) in polyQ80-GFP-v2A-GFP-injected +/+ embryos at 48hpf after treatment with Rapa or CQ. Means with standard deviations (SDs) are shown. (C) Western immunoblots from polyQ80-GFP-v2A-GFP-injected +/+ embryos at 24 hpf, 48 hpf, 72 hpf, and 96 hpf. (D) Ratios (polyQ80-GFP / free GFP) in polyQ80-GFP-v2A-GFP-injected +/+ embryos at 24 hpf, 48 hpf, 72 hpf, and 96 hpf. Means with SDs are indicated.

An advantage of zebrafish over mammalian models is the easy treatment of their living embryos with drugs by placement of these in the embryos’ aqueous support medium for direct absorption. (The absorption of less soluble drugs can be facilitated by co-exposure to 1% DMSO, e.g. (18)). To test whether polyQ80-GFP was being degraded by autophagy selectively relative to free GFP we sought to enhance autophagy induction or block autophagic flux using rapamycin, (19) or chloroquine (20). Both drugs have previously been used successfully to modulate autophagy in zebrafish (19, 21).

In zebrafish embryos, autophagy is up-regulated at the pharyngula stage (24~48 hpf), and the earliest time point at which the phosphatidylethanolamin (PE)-Lc3-II conjugate (critical for autophagy induction) has been detected is 32 hpf (22). Therefore, in our research, injected embryos were exposed to 1μM rapamycin or 50mM chloroquine from 30 hpf before lysis at 48 hpf for analysis. The ratios of polyQ80-GFP to free GFP observed from both drug treatments compared to non-treatment controls are presented in Fig 3B. The ratio (poly80Q-GFP / free GFP) was significantly reduced through rapamycin treatment (*p*=0.0090), indicating that autophagy was induced. The ratio (poly80Q-GFP / free GFP) was apparently increased through chloroquine treatment (*p*=0.0708), consistent with inhibition of autophagy. Since the changes in the ratios of poly80Q-GFP / free GFP are consistent with the changes in autophagy expected from these different drug treatments, the polyQ80-GFP-v2A-GFP assay appears able to measure autophagic flux in zebrafish.

The polyQ80-GFP / free GFP ratio observed from the polyQ80-GFP-v2A-GFP transgene possibly decreases between 24 hpf and 96 hpf as indicated by a statistically non-significant trend (Fig 3C and 3D) Also, we noticed that this assay itself is somewhat toxic to zebrafish embryos with approximately half of the injected embryos showing abnormal development at 24 hpf (data not shown). This may be due to the previously observed toxicity of polyglutamine proteins in zebrafish (23, 24). Furthermore, in 2007, Schiffer et al. reported that polyQ102-GFP could aggregate to form large SDS-insoluble inclusions, while free GFP was observed to be produced by removal of polyQ moieties from polyQ-GFP fusion proteins (23). All these phenomena might introduce unanticipated variability into observed polyQ80 / free GFP ratios (although this can be overcome somewhat by the extensive experimental replication that is facilitated by use of the zebrafish model system). For this reason we sought a less-toxic, aggregation-independent, but still internally-controlled alternative assay system.

### GFP-LC3-GFP probe to quantify autophagic flux by western immunoblotting

In 2016, Kaizuka et al. (10) described constructs to visualize autophagic flux in zebrafish embryos. Their GFP-LC3-RFP fusion protein is cleaved into separate GFP-LC3 and RFP proteins by embryos’ endogenous ATG4 activity. The GFP-LC3 then associates with autophagosomes while the RFP acts as an internal control.

We wished to adapt the assay from Kaizuka et al. to allow quantification of autophagic flux by western immunoblotting while avoiding the difficulties posed by use of polyQ80-GFP. However, we knew from previous unpublished work with GFP-v2A-RFP fusion proteins that use of an RFP internal control for normalization of GFP expression is problematic since results are only comparable between samples on the same immunoblot. This difficulty could be overcome if GFP served both as autophagy target and as the internal control. Therefore we applied the tandem GFP principle of the polyQ80-GFP-v2A-GFP construct to create GFP-LC3-GFP. This is cleaved by ATG4 within cells to give, initially, equimolar amounts of GFP-LC3 and free GFP proteins (Fig 2B). When assayed, these proteins have discernibly different molecular masses identifiable on western immunoblots probed with a single anti-GFP primary antibody (Fig 2B). The mean GFP-Lc3a / free GFP ratio observed in wild type (+/+) embryos at 48 hpf is slightly higher than at 96 hpf (1.22 vs 1.13 in Fig 5A and 5B respectively). However, similar to the polyQ80-GFP-v2A-GFP assay, this difference is not statistically significant (*p*=0.42).

To test whether changes in the GFP-Lc3a / free GFP ratio reflect changes in autophagic flux, the GPF-Lc3a-GFP construct was expressed in embryos subjected to treatment with rapamycin or chloroquine. Treatment with rapamycin produced a significant decrease in the GFP-Lc3a / free GFP ratio (*p*=0.001, Fig 5A), reflecting the expected increase in autophagy. Unexpectedly, treatment with chloroquine, both at high (50mM) and low (50μM) concentrations resulted in significant decreases in the GFP-Lc3a / free GFP ratio (Fig 5A). This apparent increase in autophagy conflicted with chloroquine’s activity as an autophagy inhibitor, which implied that an unanticipated factor could be distorting the observed ratio during the chloroquine treatments.

**Fig 4.**
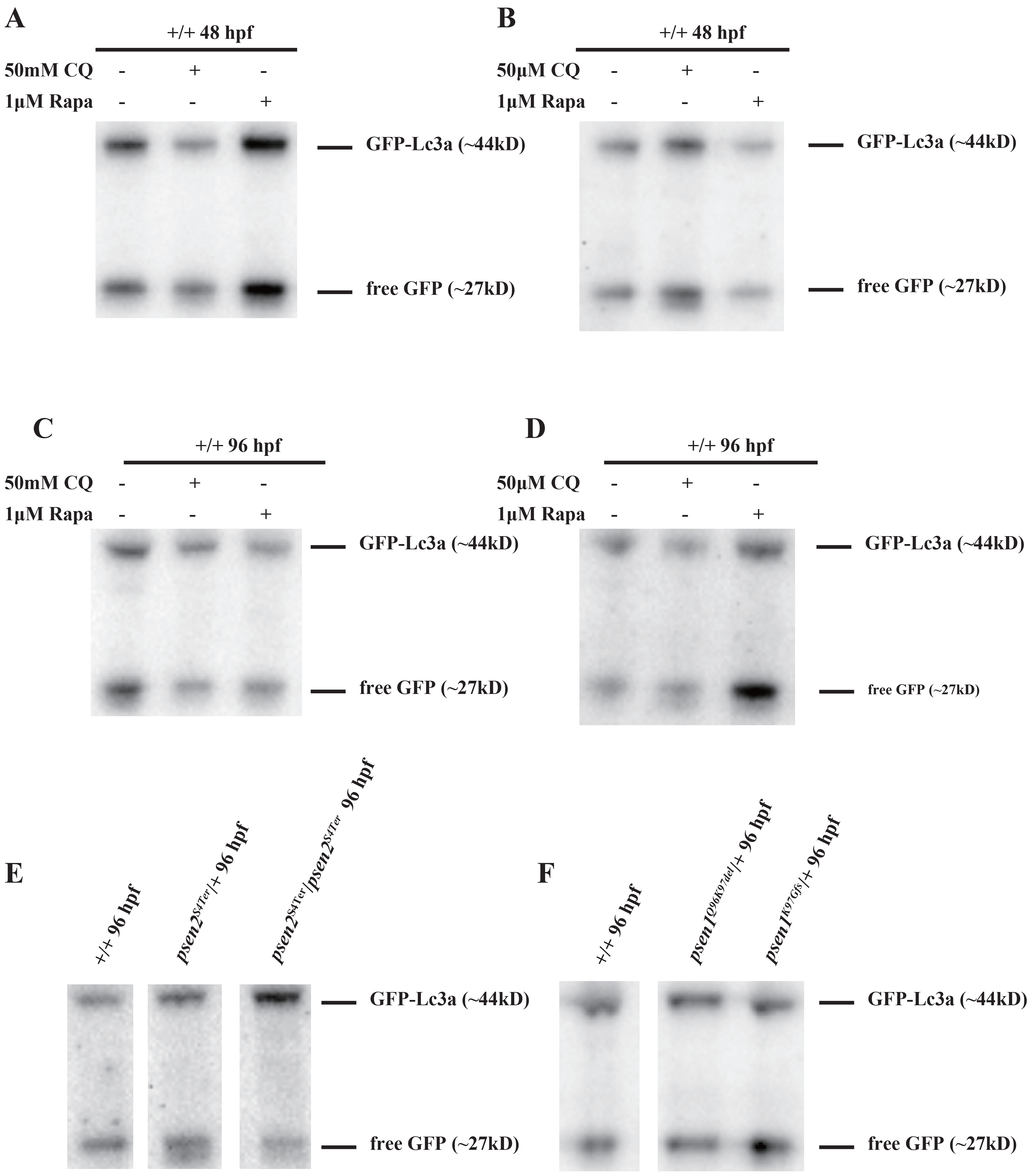

**Fig 5.**
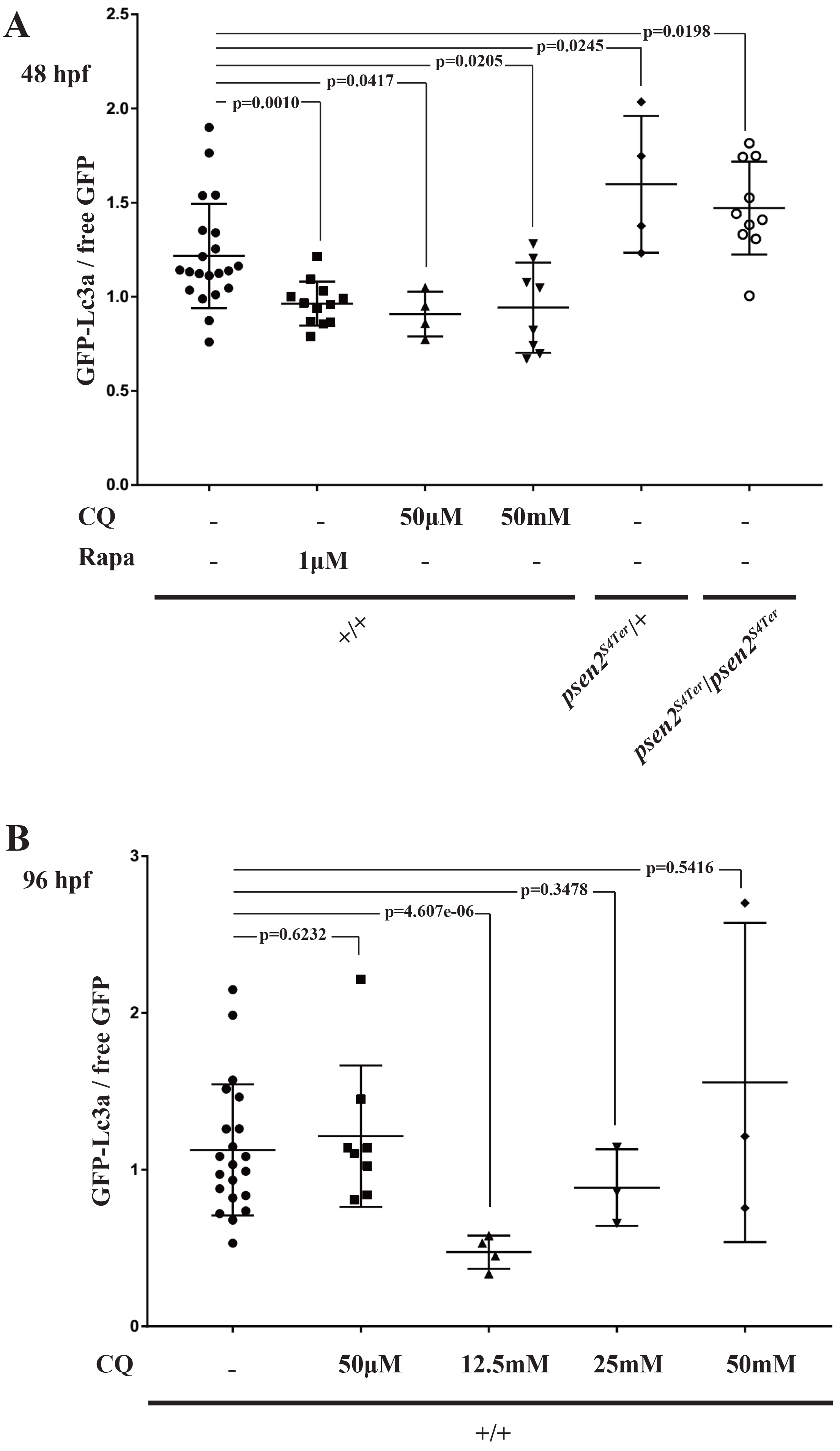
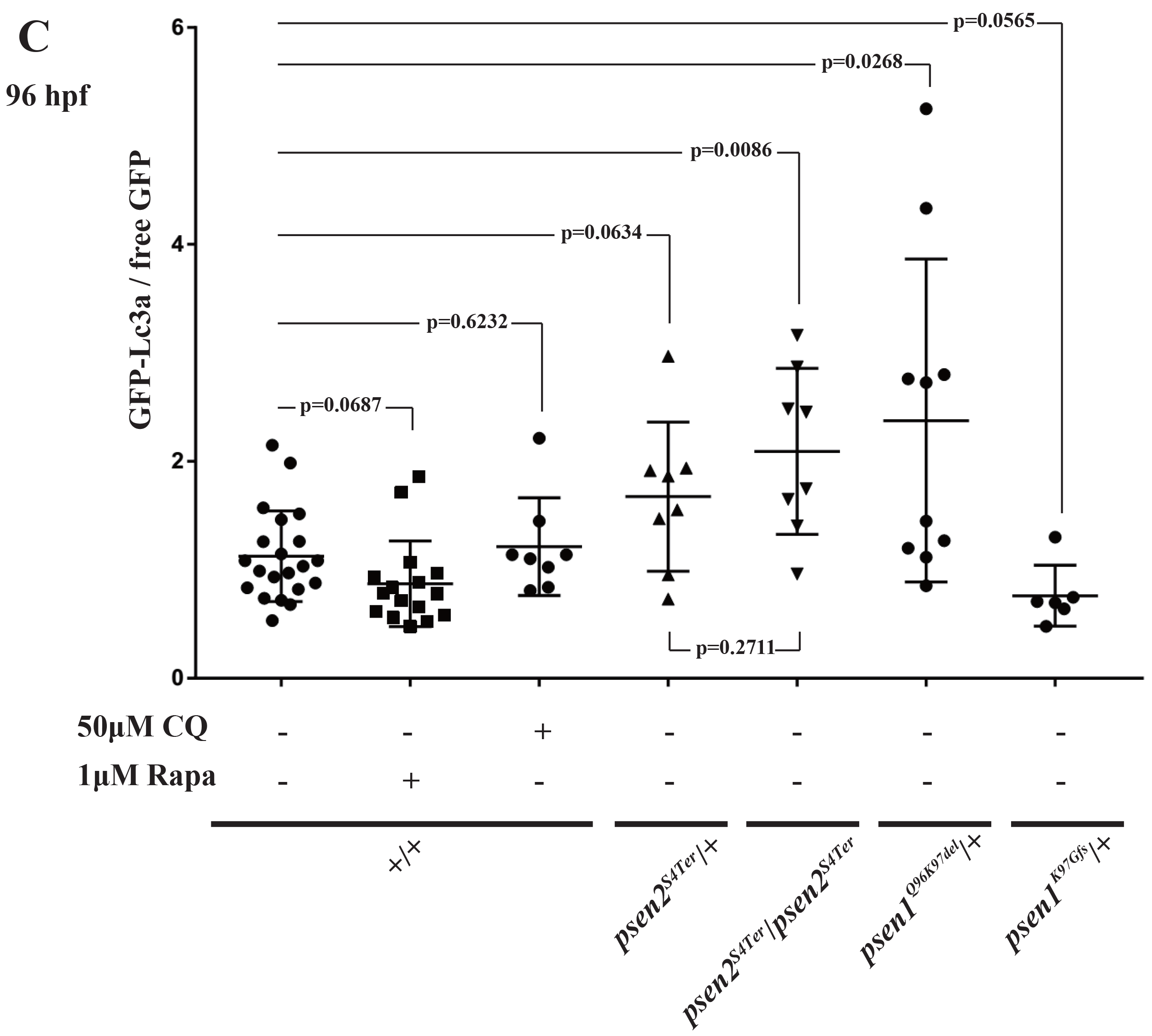
Ratios (GFP-Lc3a / free GFP) from the GFP-Lc3a-GFP assay of autophagy. A. Embryos at 48 hpf. +/+ embryos were treated with rapamycin, Rapa, or various concentrations of chloroquine, CQ. *psen2^S4Ter^* mutant embryos were not treated with these drugs.
B. +/+ embryos at 96 hpf after treatment with various concentrations of chloroquine.
C. Zebrafish embryos of various genotypes at 96 hpf. Only +/+ embryos were treated with rapamycin or chloroquine as indicated.

p-values are from two-tailed t-tests assuming either equal or unequal variances as appropriate.

After investigating the scientific literature we discovered a report from Ni et al. in 2011 describing that both chloroquine and rapamycin treatments of HeLa cells can cause increased lysosomal pH and cleavage of GFP-LC3 to release free GFP (similar to the removal of polyQ from polyQ-GFP as observed by Schiffer et al (23)). If this form of cleavage also existed in zebrafish it might provide an additional source of free GFP that would affect observed GFP-Lc3a / GFP ratios in embryos expressing the GPF-Lc3a-GFP assay transgene. To test this possibility we constructed a simple fusion of GFP to Lc3a by deleting the downstream GFP coding sequences from the GFP-Lc3a-GFP transgene to create GFP-Lc3a (Fig 2C). The GFP-Lc3a transgene was then expressed in zebrafish embryos in the same way as for GFP-Lc3a-GFP and the rapamycin and chloroquine treatments were then repeated. At 48 hpf, free GFP (i.e. not fused to Lc3a) can clearly be observed on western immunoblots (Fig 6A), confirming that a protease activity that separates GFP from Lc3a exists in zebrafish. Interestingly, the GFP-Lc3a protein from this transgene is apparently observable as both GFP-Lc3a-I and GFP-Lc3a-II forms whereas we have only observed a single form of GFP-Lc3a protein when this is produced from the GFP-Lc3a-GFP transgene. A similar observation was made by Kaizuka et al. (10) for expression of GFP-LC3 from their GFP-LC3-RFP-LC3ΔG construct when this was expressed from a transgene (as our protein fusions are) rather than from injected mRNA. The free-GFP derived from our GFP-Lc3a transgene is certainly non-negligible relative to the observed GFP-Lc3a-I (e.g. in untreated, wild type embryos, see Fig 6A) at 48 hpf. This implies that free GFP from cleavage of GFP-Lc3a may significantly affect GFP-Lc3a / GFP ratios from GFP-Lc3a-GFP-injected embryos at this developmental time point. When we observed cleavage of GFP-Lc3a in 96 hpf larvae, far less free GFP was seen relative to GFP-Lc3a-I. Therefore, the ratio of GFP-Lc3a to GFP may be less distorted by free GFP from cleavage of GFP-Lc3a at 96 hpf.

**Fig 6.**
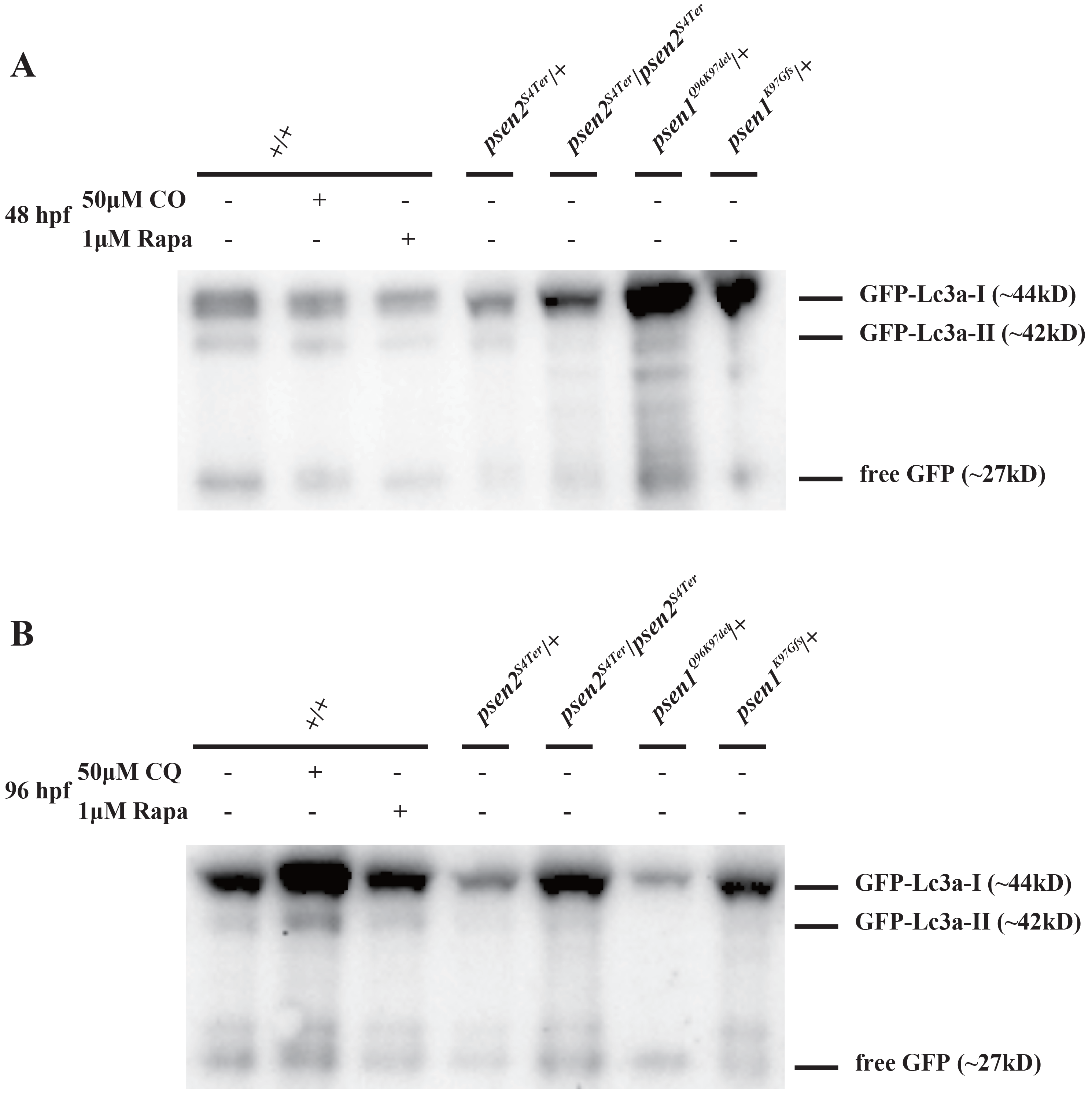
Western immunoblots from GFP-Lc3a-injected embryos and larvae. (A) 48 hpf and (B) 96 hpf respectively. +/+ embryos and larvae were also treated with rapamycin (Rapa) or chloroquine (CQ) as indicated.

When we repeated the chloroquine treatments on GFP-Lc3a-GFP transgene-injected zebrafish embryos with lysis at 96hpf we then observed a slightly increased GFP-Lc3a / free GFP ratio at the lowest chloroquine dosage (50μM) although without apparent statistical significance (*p*=0.6232, Fig 5B). At higher concentrations of chloroquine (12.5 mM and above) the GFP-Lc3a / free GFP ratio was decreased or was very variable (Fig 5B). Ni et al. reported that saturating doses of chloroquine in HeLa cells are able to block GFP-LC3 cleavage completely (25) and this may also be occurring in some zebrafish embryos treated with 50 mM chloroquine. A large proportion of larvae (~80%) were dead at 96 hpf after the 50mM chloroquine treatment indicating high lethality of this chloroquine concentration.

We conclude that exploitation of the GFP-Lc3a-GFP assay is best performed at 96hpf or later to minimize the influence of generation of free GFP from cleavage of GFP-Lc3a. Generation of free GFP from cleavage of GFP-Lc3a reduces the sensitivity of the GPF-Lc3a-GFP construct for detecting decreases in autophagic flux.

### Autophagic flux in zebrafish *psen* mutants expressing GFP-Lc3a-GFP

Our tests of the GFP-Lc3a-GFP assay indicated that, when used at 96 hpf, it can reveal changes in autophagic flux without the toxicity and other problems encountered with the aggregation-based polyQ80-GFP-v2A-GFP assay. Therefore, we exploited the GFP-Lc3a-GFP assay to examine autophagic flux changes in a number of zebrafish *presenilin* mutants that we have generated in our laboratory (S3 File). These are: a typical fAD-like mutation in *psen1* lacking two codons but preserving the open reading frame, *psen1^Q96K97del^*; a putatively null mutation of the *psen2* gene, *psen2^S4Ter^*; and a zebrafish model of the unique, open reading frame-truncating PSEN fAD mutation *PSEN2^K115Efs^* (14). We have previously suggested that this latter mutation causes fAD by inappropriate mimicry of expression of an hypoxia-induced, naturally-occurring truncated isoform of PSEN2 denoted PS2V (26). In zebrafish, the PS2V-equivalent isoform is expressed from the *PSEN1*-orthologous gene, *psen1*. Therefore, we have modeled this mutation in zebrafish by generation of the *psen1* allele, *K97Gfs*.

The effects of the fAD-like mutations were tested in the heterozygous state corresponding to the dominant inheritance pattern shown by such mutations in humans. The effects of the putatively null mutation of *psen2* were tested in both homozygous and heterozygous embryos to examine, respectively, whether *psen2* activity has any effect on autophagy and whether this effect is haploinsufficient. Fertilized eggs bearing *presenilin* mutations were injected with the GFP-Lc3a-GFP transgene and then examined by western immunoblotting at 96 hpf (Fig 5C). Both heterozygosity for the fAD-like *psen1^Q96K97del^* mutation and loss of *psen2* activity were observed to decrease autophagic flux significantly (*p*=0.0268 and *p*=0.0086 respectively) as expected from previous analysis of fAD mutations in human *PSEN1* (8) and confirming that *psen2* also functions in autophagy. Heterozygosity for the putatively null mutation in *psen2* also appeared to reduce the GFP-Lc3a / free GFP ratio indicating decreased autophagic flux although this did not reach statistical significance (*p*=0.063). Unexpectedly, heterozygosity for the mutation *psen1^K97Gfs^* that models the unique ORF-truncating *K115E* fAD mutation of human *PSEN2* appeared possibly to increase autophagic flux (*p*=0.0565) reflecting complexity in the relationship between fAD mutations and this crucial cellular process.

## Discussion

To monitor autophagic flux in zebrafish embryos, we first designed the polyQ80-GFP-v2A-GFP assay that is based on the assumption that polyQ80-GFP aggregates and is a target for autophagy while free GFP is primarily degraded via the proteasome. Since this assay provides 1:1 stoichiometric co-expression of polyQ80-GFP and free GFP protein within the same cells *in vivo* it should reflect changes in autophagic flux by changes in the ratio of polyQ80-GFP / free GFP as observed by western immunoblotting. Since both the polyQ80-GFP and the free GFP protein are detected simultaneously by the same primary antibody, polyQ80-GFP / free GFP ratios can be compared on separate immunoblots which greatly facilitates experimental replication and statistical analysis. While rapamycin and chloroquine treatments of embryos injected with the polyQ80-GFP-v2A-GFP transgene produced ratio changes indicative of the expected changes in autophagic flux, the aggregation of polyQ80 may, itself, induce autophagy and this would limit the assay’s sensitivity. This issue is overcome by replacing the polyQ80-GFP protein with a non-aggregating GFP-Lc3a fusion protein, based on the approach taken by Kaizuka et al. (10). Like the polyQ80-GFP-v2A-GFP construct, the GFP-Lc3a-GFP construct produces 1:1 stoichiometric co-expression of two proteins in the same cells *in vivo* (GFP-Lc3a and free GFP proteins). The GFP-Lc3a-GFP assay assumes that GFP-Lc3a protein is a target of autophagy while the free GFP protein remains in the cytosol as an internal control. However, the GFP-Lc3a-GFP transgene should not affect autophagic flux itself nor TORC1 activity as the related GFP-LC3-RFP-LC3ΔG construct of Kaizuka et al. did not appear to do so (10). Nevertheless, the GFP-Lc3a-GFP assay has its own particular limitations. Cleavage of LC3 away from GFP was reported in HeLa cells when lysosomal pH was increased by chloroquine or rapamycin treatments (25). Consistent with this, we also saw production of free GFP by cleavage of GFP-Lc3a in zebrafish embryos and larvae (Fig 6A and 5B). Since the free GFP produced by GFP-Lc3a cleavage provided an additional source of free GFP, this can affect the measured ratio of GFP-Lc3a / free GFP and this reduces the sensitivity of the assay to detect decreases in autophagic flux. We found that this effect was minimized at 96 hpf compared to 48 hpf. (Note that the observations by Schiffer et al. imply that polyQ80-GFP will also form free GFP by removal of polyQ80 in zebrafish embryos (23).)

It is possible that development of a GFP-Lc3a-RFP-GFP transgene-based autophagy assay might overcome the insensitivity caused by generation of free GFP from GFP-Lc3a since an RFP-GFP fusion would be easy to identify separately from any free GFP in western immunoblotting. The accumulation of free GFP versus GFP-Lc3a in such an assay would be additionally informative regarding autophagic flux versus lysosomal accumulation.

Using the GFP-Lc3a-GFP assay, we were able to compare the levels of autophagic flux in zebrafish mutant and wild type larvae. In both *psen2^S4Ter^*/+ and *psen2^S4Ter^*/*psen2^S4Ter^* larvae, we detected decreased autophagic flux (Fig 5C), demonstrating that *psen2*, like *psen1*, plays a role in regulating cells’ autophagic activity. This has not previously been shown. However, we should note that, in terms of Psen2 protein’s role in γ-secretase activity, our previous work has shown that zebrafish Psen2 plays a greater role in Notch signaling than its corresponding mammalian orthologue (27) and we cannot exclude that zebrafish Psen2 is more similar to Psen1 in its role in autophagy than mammalian PSEN2 is similar to PSEN1. In heterozygous larvae of the “typical” (reading frame-preserving) fAD mutation model *psen1^Q96K97del^*, we also observed decreased autophagic flux (Fig 5C). Interestingly, our zebrafish model of the human *PSEN2^K115Efs^* fAD mutation, *psen1^K97Gfs^*, did not show decreased autophagic flux. Instead, heterozygosity for this mutation appeared possibly to increase autophagic flux in the GFP-Lc3a-GFP assay.

The *K115Efs* mutation of human *PSEN2* is unique among the human *PRESENILIN* fAD mutations in that it truncates the open reading frame of the gene. Nevertheless, our unpublished analyses of aged adult fish modelling this mutation show similar alterations in expression of the genes responding to hypoxia to those seen in aged fish carrying the more “typical” fAD-like mutation *Q96K97del* (Newman et al. *manuscript in preparation*). Therefore, it will be interesting to examine autophagic flux in such aged fish to see whether it subsequently becomes inhibited (as is typical in Alzheimer’s disease (4)). To this end, we are currently determining whether the GFP-Lc3a-GFP transgene can be transmitted through the zebrafish germline for widespread expression in adult tissues.

The possibly increased autophagic flux observed due to heterozygosity for the reading frame-truncating *psen1^K97Gfs^* mutation argues against the idea that the human *PSEN2^K115Efs^* mutation, or the zebrafish *psen1^K97Gfs^* model of it, represent hypomorphic alleles as such alleles would be expected to decrease autophagic flux (e.g. see (8) and the putatively *psen2* null allele analysis of this paper). In fact, our results suggest that this truncated form of PRESENILIN, and the naturally occurring PS2V isoform that it supposedly mimics, may interact with PRESENILIN holoproteins to increase their function in autophagy. Induction of autophagy is a pro-survival response to cellular stresses such as hypoxia (28) and we have previously shown using zebrafish that PS2V acts to decrease cell death under such conditions (26). The observation of increased autophagic flux in zebrafish larvae expressing *psen1^K97Gfs^* is consistent with this.

In summary, we tested two GFP-based assays intended to detect changes in autophagic flux in zebrafish embryos and larvae by western immunoblotting: polyQ80-GFP-v2A-GFP and GFP-Lc3a-GFP. Both transgenes provide 1:1 stoichiometric co-expression of an autophagy target protein and of free GFP as an internal control. The two assays both appeared able to reflect changes in autophagic flux although each assay displayed particular limitations. Using the GFP-Lc3a-GFP assay, we found that lack of *psen2* activity in zebrafish reduces autophagic flux as does heterozygosity for a typically reading-frame-preserving fAD-like mutation of *psen1* (*Q96K97del*).

## Supporting information

**S1 File. Sequence design and sub-cloning for the GFP-based constructs.**

(A) Sequence design for the polyQ80-GFP-v2A-GFP construct

(B) Sequence design for the GFP-Lc3a-GFP construct

(C) Sequence design for the GFP-Lc3a construct

(D) Sub-cloning of the GFP-based constructs into the Tol2 vector

**S2 File. Intensity ratios of western immunoblots**

Table 1. Intensity ratios of western immunoblots for Figure 3B

Table 2. Intensity ratios of western immunoblots for Figure 3D

Table 3. Intensity ratios of western immunoblots for Figure 5A

Table 4. Intensity ratios of western immunoblots for Figure 5B

Table 5. Intensity ratios of western immunoblots for Figure 5C

**S3 File. Mutagenesis and breeding of the mutant fish**

(A) Mutagenesis and breeding of the *psen2^S4Ter^* zebrafish

(B) Mutagenesis and breeding of the *psen1^Q96K97del^* and *psen1^K97Gfs^* zebrafish

## Abbreviations

Aβ: amyloid-β
AD: Alzheimer’s Disease
APP: Amyloid beta A4 protein
Atg: autophagy-related gene
AV: autophagic vacuole
CQ: chloroquine
GFP: green fluorescent protein
LC3: microtubule-associated protein 1 light chain 3
PE: phosphatidylethanolamin
polyQ: poly-glutamine
PSEN: presenilin; Rapa, rapamycin

